# Direct quantification of unicellular algae sinking velocities reveals cell size, light, and nutrient-dependence

**DOI:** 10.1101/2023.06.20.545838

**Authors:** Teemu P. Miettinen, Annika L. Gomez, Yanqi Wu, Weida Wu, Thomas R. Usherwood, Yejin Hwang, Benjamin R.K. Roller, Martin F. Polz, Scott R. Manalis

## Abstract

Eukaryotic phytoplankton, also known as algae, form the basis of marine food webs and drive marine carbon sequestration when their biomass sinks to the ocean floor. Algae must regulate their vertical movement, as determined by motility and gravitational sinking, to balance access to light at the surface and nutrients in deeper layers. However, the regulation of gravitational sinking velocities remains largely unknown, especially in motile species. Here, we directly quantify single-cell masses and volumes to calculate sinking velocities according to Stokes’ law in diverse clades of unicellular marine microalgae. Our results reveal the cell size, light, and nutrient-dependency of sinking velocities. We identify motile dinoflagellate and green algal species that increase their sinking velocity in response to starvation. Mechanistically, this increased cell sinking is achieved by photosynthesis-driven accumulation of carbohydrates, which increases cell mass and density. Moreover, cell sinking velocities correlate inversely with proliferation rates, and the mechanism regulating cell sinking velocities integrates signals from multiple nutrients. Our findings suggest that the regulation of cell composition according to environmental conditions contributes to the vertical movement of motile cells in the oceans. More broadly, our approach for sinking velocity measurements expands the study of gravitational sinking to motile cells and supports the modeling of marine carbon pump and nutrient cycles.

## INTRODUCTION

Algae are a genetically, morphologically, and biophysically diverse group of photosynthetic microorganisms. Algae comprise the base of the marine food webs and are responsible for approximately 75-90% of marine carbon fixation by mass ^1^. Consequently, the sinking of algal material – either in the form of single cells or as components of larger sinking particles, such as marine snow ^2–4^ – is a key contributor to the ‘biological pump’, which sequesters carbon dioxide from the atmosphere to deep oceans and marine sediments ^5^. Hence, a quantitative understanding of sinking velocities of different algae is needed for modeling marine ecosystems and carbon fluxes, especially in the context of global change ^2,6–11^.

The vertical position of live cells impacts their growth, viability, and community composition, as both organic and inorganic nutrients are more abundant deeper in the ocean while light is stronger at the surface ^7,12^. Hence, algae are under ecological pressure to adjust their position in the water column according to their need for nutrients and light. To adjust their vertical position, algae can alter their gravitational sinking and motility, but live cell sinking and sedimentation rate measurements cannot decouple gravitational sinking from motility. Consequently, studies on gravitational sinking have focused on non-motile species, which can adjust their gravitational sinking by regulating cell size and composition ^13–17^. The extent to which gravitational sinking is regulated in motile species remains unclear, and the molecular mechanism(s) responsible for such sinking regulation are unknown even in most non-motile species.

Gravitational sinking velocity depends on the cell’s size, density and drag force from seawater. For spherical cells, this can be derived from the Stokes’ law ^3,6,13,14^:

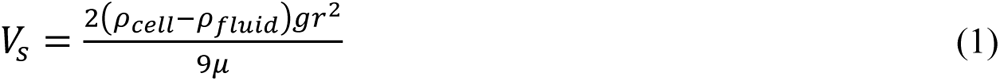

where *p_cell_* is the density of the cell, *p_fluid_* is the density of the surrounding fluid, *g* is the gravitational acceleration, *r* is radius of the cell, and *μ* is the dynamic viscosity of the fluid. A cell dimension-dependent correction factor can be applied to *Eq 1* for non-spherical particles ^14,18^. This approach has been widely used to estimate sinking velocities based on microscopy observations of cell radius, but cell density is rarely known or directly measured, especially for single cells. Consequently, little is known about the cell size-dependency of sinking velocities on a single-cell level.

We reasoned that independent measurements of cell volume and buoyant mass could be used to calculate sinking velocity without assumptions about cell density (Fig. 1A). Buoyant mass, *m_b_*, is defined as

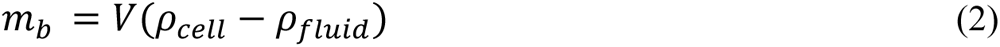

**Fig. 1.**
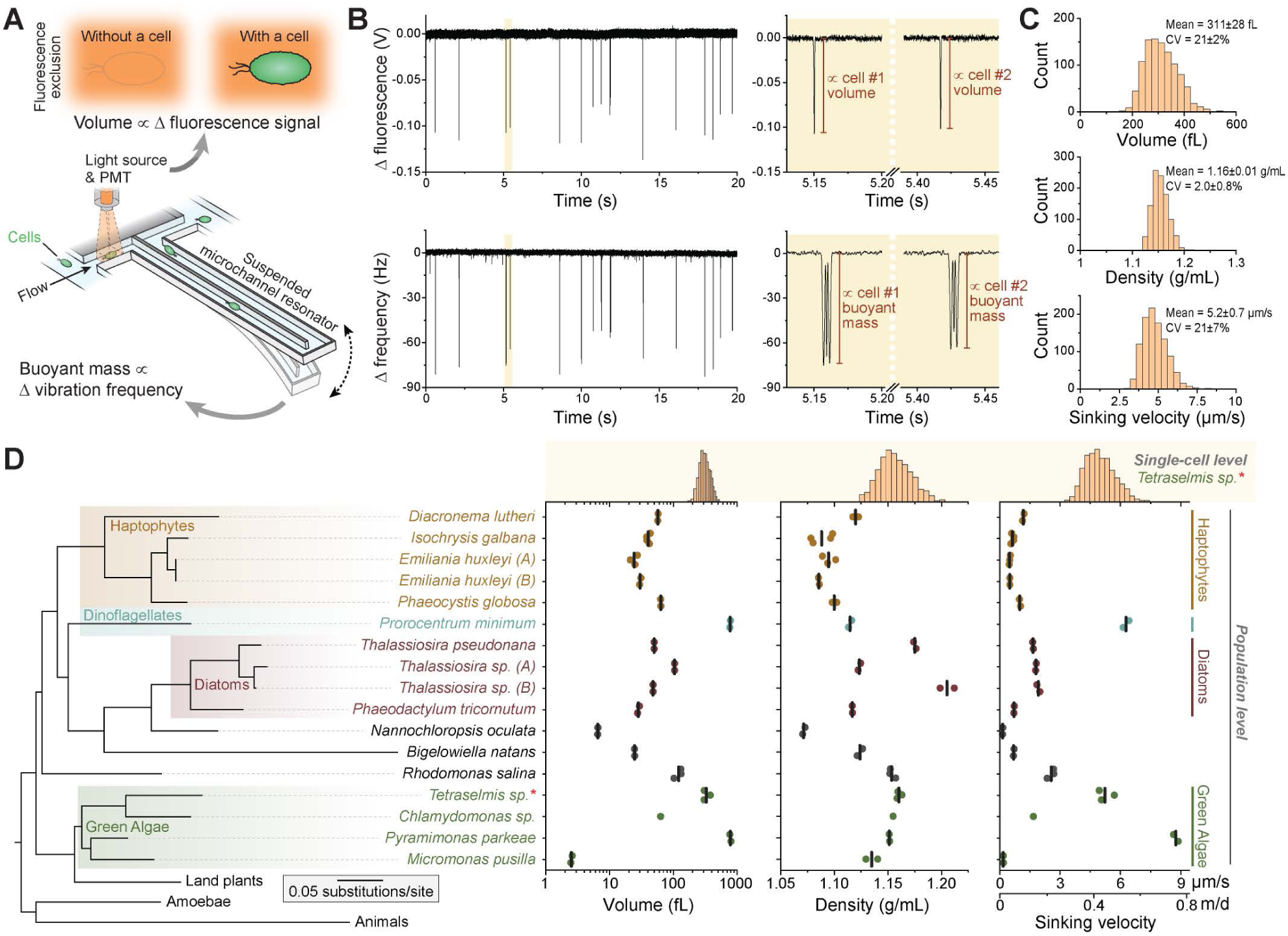
Measuring gravitational sinking velocities across phylogenetically diverse algae species and single cells. **(A)** Schematic of the measurement approach. To measure gravitational sinking velocities according to Stokes’ law on a single-cell level, we coupled the suspended microchannel resonator-based buoyant mass measurements with fluorescence exclusion-based volume measurements. **(B)** *Left,* representative raw data for fluorescence exclusion measurements (top) and buoyant mass measurements (bottom) for the species *Tetraselmis sp*. *Right*, zoom-ins displaying data for two example cells. **(C)** Representative single-cell volume, density, and sinking velocity histograms for *Tetraselmis sp*. (n=1027 single cells). The listed statistics depict mean±SD from 5 independent cultures. **(D)** *Left*, a phylogenetic tree of the measured algae species according to 18S rRNA. Animals, amoebae, and land plants are shown for reference. The major clades of algae are color coded. *Right*, cell volumes, densities, and sinking velocities for each species. Dots depict independent cultures and vertical lines depict mean values. *Top,* Representative histograms of single-cell volumes, densities and sinking velocities for the species *Tetraselmis sp*.

where, *V* is the volume of the cell. As the fluidic density is typically known, cell density, and consequently sinking velocity, can be found from buoyant mass and volume. Here, we obtain direct measurements of single-cell volumes and buoyant masses to define algae sinking velocities across different species, cell sizes, and light and nutrient conditions. Our work reveals examples of motile species that increase their sinking velocity upon starvation, and we uncover the mechanistic basis behind this sinking velocity regulation. Overall, our results allow us to propose a qualitative ecological model where the regulation of cell size and composition impacts marine carbon pump and nutrient cycles.

## RESULTS

### Characterizing sinking velocities across phylogenetically diverse algae species and single cells

We sought to establish an approach for measuring gravitational sinking velocities across algal species and across single cells within a species. We coupled the suspended microchannel resonator (SMR) ^19–22^ with fluorescence exclusion-based volume measurements ^23,24^ (Fig. 1A). The SMR is a microfluidic mass sensor, where cells flow through a microchannel that is embedded in a vibrating cantilever. This changes the vibration frequency of the cantilever proportionally to the buoyant mass of the cell (Fig. 1B). Next to the SMR cantilever, our system incorporates a fluorescence detection area, where the fluorescence signal decreases proportionally to the volume of a cell travelling through, as the cell excludes cell-impermeable fluorophore-containing media from the detection area (Fig. 1B). Our SMR mass precision is ∼0.1% ^20,22^, and our volume precision by fluorescence exclusion is ∼1% ^23^. Thus, we can precisely determine single-cell volume and buoyant mass, which we use to calculate cell density and sinking velocity for individual algal cells (*Eq 1&2*).

To the best of our knowledge, single-cell densities and gravitational sinking velocities have never been reported for motile algae. We characterized two unicellular eukaryotic algal species: *Tetraselmis sp.*, which is an industrially relevant green alga with high lipid production capacity and uses in aquaculture ^25,26^, and *Prorocentrum minimum*, which is a globally distributed, bloom-forming dinoflagellate that produces toxins impacting shellfish populations and human health ^27^. The single-cell densities of *Tetraselmis sp.* displayed a coefficient of variation (CV) of 2.0 ± 0.8% (mean ± SD) (Fig. 1C), which is an order of magnitude more than what is observed in typical eukaryotic model systems ^28^. Consequently, there was significant cell-to-cell variability in sinking velocities with CV values of 21 ± 7% (mean ± SD). *Prorocentrum minimum* displayed even higher variability in single-cell densities (CV of 2.7 ± 0.7%, mean ± SD) and sinking velocities (CV of 38 ± 9%, mean ± SD) than *Tetraselmis sp.* (Supplemental Fig. S1A). Thus, cell densities and sinking velocities in algae are more variable than expected based on observations in other model systems.

In addition to our single-cell sinking velocity measurements, we used a population level approach to examine a range of unicellular algal species. Using the SMR and Coulter counter to measure buoyant masses and cell volumes, respectively, we characterized cell densities and sinking velocities in 17 unicellular eukaryotic algal species spanning all major clades of algae, including haptophytes, dinoflagellates, diatoms, and green algae (Fig. 1D, *left*, Supplemental Fig. S1B). All species were verified using 18S rRNA sequencing and grown under autotrophic conditions in high nutrient media. The measured species spanned ∼2.5 orders of magnitude in volume from the ∼2.5 fL green alga *Micromonas pusilla* to the ∼785 fL green alga *Pyramimonas parkeae* (Fig. 1D, Supplemental datasets S1&S2). Despite the wide phylogenetic and cell volume range of the studied species, cell densities ranged only ∼12% between the species. We also observed significant density variation between two separate isolates of the diatom *Thalassiosira sp.* (Fig. 1D), thus demonstrating the degree of density variation possible among closely related taxa. Cell sinking velocities ranged ∼410% between species, from 0.2 µm/s in *Nannochloropsis oculata* to 12.8 µm/s in *Pyramimonas parkeae*. Notably, the cell shape-dependency of sinking velocities was minimal (Supplemental dataset S3), and all sinking velocities reported in figures therefore assume a spherical cell shape. Overall, these results reveal the interspecies heterogeneity in cell densities and sinking velocities and indicate that our measurement approach is suitable for a wide size and phylogenetic range of algal species.

In oceans, individual algae species are often dispersed across wide vertical distances. Similarly, the sinking of individual cells may not be well represented by the mean sinking velocity of the population. To provide context for our single-cell observations of cell density and sinking velocity, we compared the cell-to-cell variation observed in *Tetraselmis sp.* to the species-to-species variation. The density variation within a species (CV ∼2%) is similar in magnitude to the density variation between species (CV ∼3%) (Fig. 1D). Single-cell sinking velocities also varied across a wide part of the sinking velocity range observed across species (Fig. 1D, *top*). These results suggest that, in addition to active motility and cell aggregation, the variability of cell sinking velocities could result in algal dispersal across wide vertical ranges in oceans.

### ⅔-power law scaling of sinking velocities with cell volume

Understanding the cell volume-dependency of cell sinking is critical for ecological modeling, especially as cell sizes are predicted to decrease with warming oceans ^29,30^. Stokes’ law predicts that cell sinking velocities scale with cell volumes according to the ⅔-power law (*Eq 1*), assuming cell volume-independent densities. However, cell densities have not been systematically measured in most algae, making it unclear whether density is uniform across different volumes. Previous studies have suggested that algae sinking velocities scale with cell volume with a scaling factor that is significantly less than ⅔, implying that density may decrease with increasing cell volume or that cell shapes may be highly volume-dependent ^31,32^. We used our dataset (Fig. 1D) to examine the volume-dependency of cell densities and sinking velocities across species using regression models with and without phylogenetical information. In both models, cell volumes did not correlate with cell densities (Fig. 2A), but cell volumes displayed a strong correlation with sinking velocities (Fig. 2B). Over 90% of the sinking velocity variation was accounted for by cell volumes (R^2^ = 0.92-0.95). The sinking velocities scaled with cell volumes with a scaling exponent of ∼ 0.73 ± 0.04 (mean ± SE). Overall, both regression models displayed comparable results, indicating that cell volume, density, and sinking velocity have little phylogenetic signal.

**Fig. 2.**
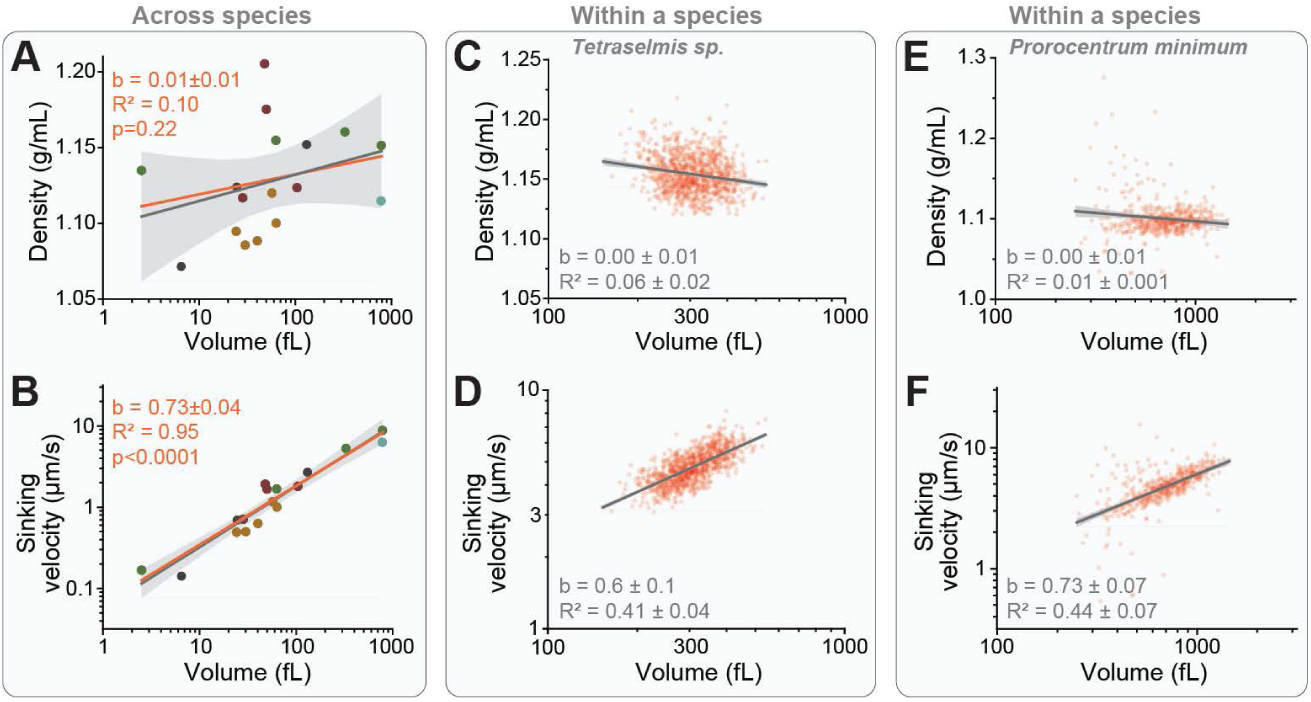
Sinking velocities display ⅔-power law scaling with cell volumes across and within species. (A-B) Correlation between cell volumes and cell densities (A) or sinking velocities (B) across all species shown in Fig. 1D. Dots depict mean values for each species (N=17), which are color coded according to their major clade. Data was fitted with linear regression models with (orange) and without (grey) phylogenetical information, and shaded areas indicate 95% confidence bands for the latter model. **(C-D)** Representative experiment displaying the correlation between cell volumes and cell densities (C) or sinking velocities (D) for individual *Tetraselmis sp*. cells (red dots, n=1027 cells). Grey lines and shaded areas indicate linear regressions and 95% confidence bands, respectively. The listed statistics were calculated from 5 separate experiments. **(E-F)** Same as (C-D), but data is for species *Prorocentrum minimum* (n=484 cells).

Our single-cell measurements also enabled us to examine the volume dependency of cell densities and sinking velocities within a species. In *Tetraselmis sp*. and *Prorocentrum minimum*, cell densities were not correlated with cell volumes (Fig. 2C,E), but cell volumes displayed a modest correlation with sinking velocities (R^2^ ∼0.4-0.5, Fig. 2D,F). Single-cell sinking velocities scaled with cell volumes with a scaling exponent of 0.6-0.7 (Fig. 2D,F). Similar results were obtained when cells were grown under low nutrient conditions (Supplemental dataset S4-5).

Overall, these results show that cell densities are not cell volume dependent, resulting in approximately ⅔-power law scaling of sinking velocities both within and between species. Consequently, when comparing vastly different sized species, cell volume can be used as a proxy for sinking velocities. However, when comparing algal cells with similar volumes, cell density differences have a more prominent impact on the sinking velocities and need to be considered. Next, we explore situations where cell sinking velocities change due to cell density changes.

### Nutrient limitation increases sinking velocity by increasing cell density in a species-specific manner

Cell volumes are known to depend on environmental nutrient conditions with higher nutrient conditions typically supporting higher cell volumes ^33^. However, little is known about how cell densities, or sinking velocities, change with environmental nutrient conditions. We therefore examined the nutrient-dependency of sinking velocities. We compared 4 algae species following 5-day culture under high and low nutrient conditions. The species investigated were *Bigelowiella natans* (chlorarachniophyte), *Phaeodactylum tricornutum* (diatom), *Tetraselmis sp.* and *Prorocentrum minimum*. The species differed in their nutrient responses, with most pronounced responses observed in *Tetraselmis sp.* (Fig. 3A) and *Prorocentrum minimum,* both of which increased their masses under low nutrient conditions while maintaining nearly constant cell volumes (Fig. 3B). Consequently, *Tetraselmis sp.* and *Prorocentrum minimum* displayed significantly increased cell densities and sinking velocities under low nutrient conditions (Fig. 3C&D). We did not observe any systematic morphological differences between the two nutrient conditions in *Tetraselmis sp.* (Fig. 3A), despite the low nutrient conditions resulting in a stationary growth phase (starvation) (Supplemental Fig. S2A). Thus, our results highlight the need for direct cell mass (or density) measurements when studying cell sinking regulation. More broadly, these results show that cells can adjust their sinking velocities significantly upon changes in environmental nutrient conditions, likely reflecting an adaptation that allows starving algae to sink faster to deeper, nutrient richer waters. However, the sinking velocity responses to nutrient limitation were species-specific, as *Bigelowiella natans* did not display any nutrient responses despite starvation under the low nutrient conditions (Fig. 3B-D, Supplemental Fig. S2A), and *Phaeodactylum tricornutum* decreased its sinking under low nutrient conditions (Fig. 3B-D).

**Fig. 3.**
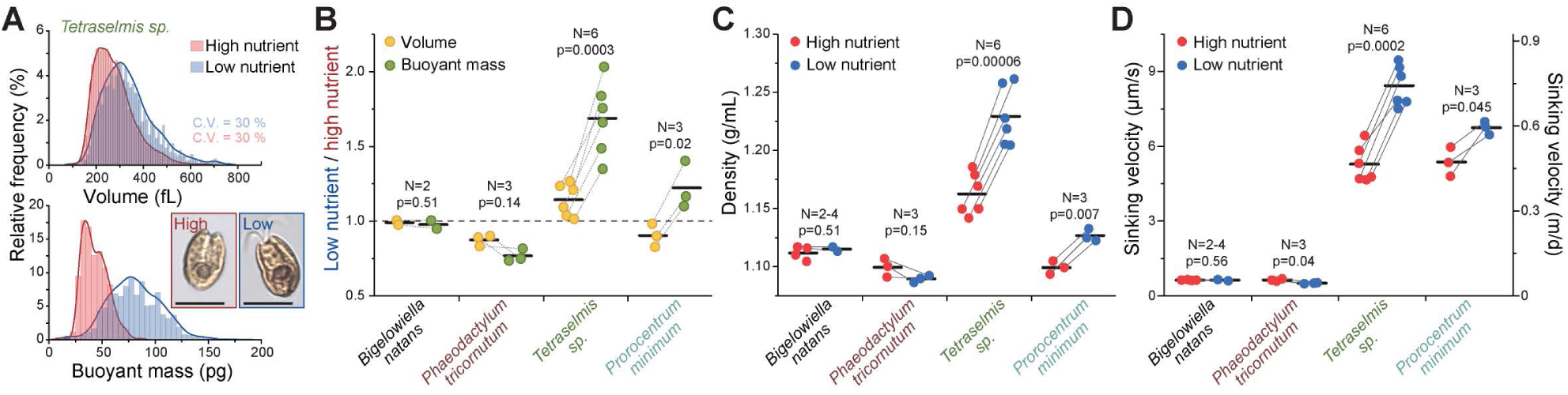
Starvation increases sinking velocity by increasing cell density in a species-specific manner. **(A)** Single-cell mass and volume histograms for *Tetraselmis sp*. in high (red) and low (blue) nutrients. Inset displays representative images of the cells. Scales bars denote 5 µm. **(B)** Change in cell volume (orange) and buoyant mass (green) between low and high nutrient conditions in indicated species. **(C-D)** Cell density (C) and sinking velocity (D) under high (red) and low (blue) nutrient conditions in indicated species. All samples were grown under high light conditions. In panels (B-D), N indicates the number of independent cultures as depicted by dots, paired samples are connected by a line, and horizontal lines depict mean values. p-values were obtained by pair-sample t-test.

### Photosynthesis-driven accumulation of carbohydrates enables sinking velocity increases

We next aimed to understand the mechanistic basis for nutrient-dependent sinking velocity regulation in more detail using *Tetraselmis sp.* as a model system. Because the increased cell densities and sinking velocities under starvation were caused by the addition of cell mass, we sought to determine if photosynthesis is required for this mass increase. We repeated our cell density and sinking velocity measurements under varying nutrient and light conditions following a 5-day culture. This revealed that the complete absence of light prevented the cells from increasing their density and sinking velocity under the low nutrient conditions (Fig. 4A-B). Measurements of cell proliferation revealed that the cells entered starvation after 2 days of low nutrient conditions but not under high nutrient conditions when light was present (Supplemental Fig. S2B). However, proliferation was stopped in the absence of light independently of nutrient availability across the time course (Supplemental Fig. S2B), as expected for a photoautotrophic species under autotrophic conditions (Fig. 4A-B). Thus, stopping cell proliferation does not automatically increase sinking velocities. More broadly, the mechanism by which cells accumulate mass to increase their density and sinking velocity requires photosynthesis.

**Fig. 4.**
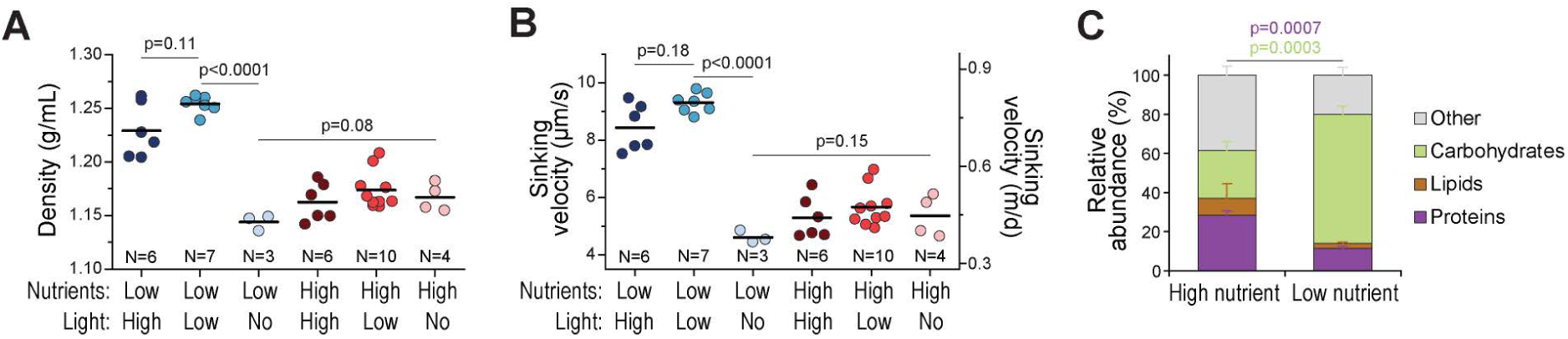
Starved *Tetraselmis sp.* increases sinking velocity by photosynthesis-driven accumulation of carbohydrates. **(A-B)** *Tetraselmis sp.* cell densities (A) and sinking velocities (B) under indicated nutrient and light conditions. p-values were obtained by ANOVA and Tukey’s posthoc test. Dots depict independent cultures and horizontal lines depict mean values, N indicates the number of independent cultures. **(C)** Relative composition of *Tetraselmis sp.* cells grown under high and low nutrient conditions. ‘Other’ category refers to ash weight. Data depicts mean±SD of 3 independent cultures. p-values for proteins (purple) and carbohydrates (green) were obtained by unpaired t-test.

Next, we studied how *Tetraselmis sp.* alter their composition to achieve the higher sinking velocity under low nutrient conditions. We characterized the protein, lipid, and carbohydrate content of the cells in relation to the cells’ dry mass. This revealed that under low nutrient conditions the cells contain less proteins, but significantly more carbohydrates, than under high nutrient conditions (Fig. 4C). Overall, these results indicate that the starvation induced cell mass and sinking velocity increases in *Tetraselmis sp.* can be explained by photosynthesis-driven accumulation of carbohydrates.

### The regulation of cell sinking velocities is responsive to multiple nutrients and coupled with the regulation of cell proliferation

Surface ocean nutrient levels vary spatially and temporally. While nitrogen is the most common growth limiting nutrient, other nutrients can also become growth limiting ^12^. To understand the nutrient-specificity of sinking velocity regulation, we first tested which nutrient is responsible for starvation and increased cell sinking under our low nutrient conditions. We aimed to rescue the starvation in the low nutrient *Tetraselmis sp.* cultures by different nutrient supplementations (Fig. 5A). While the addition of phosphorus, vitamins, or trace metals had no effect on cell growth on their own, the addition of nitrogen enabled cells partially rescued cell growth, as measured by optical density (Supplemental Fig. S2C). Direct cell counts verified the rescue of cell growth by nitrogen supplementation, although phosphorus supplementation was also capable of a minor rescue in cell counts (Fig. 5B). Together, these results indicate that nitrogen is, at least initially, the growth limiting nutrient under the low nutrient conditions used in this study.

**Fig. 5.**
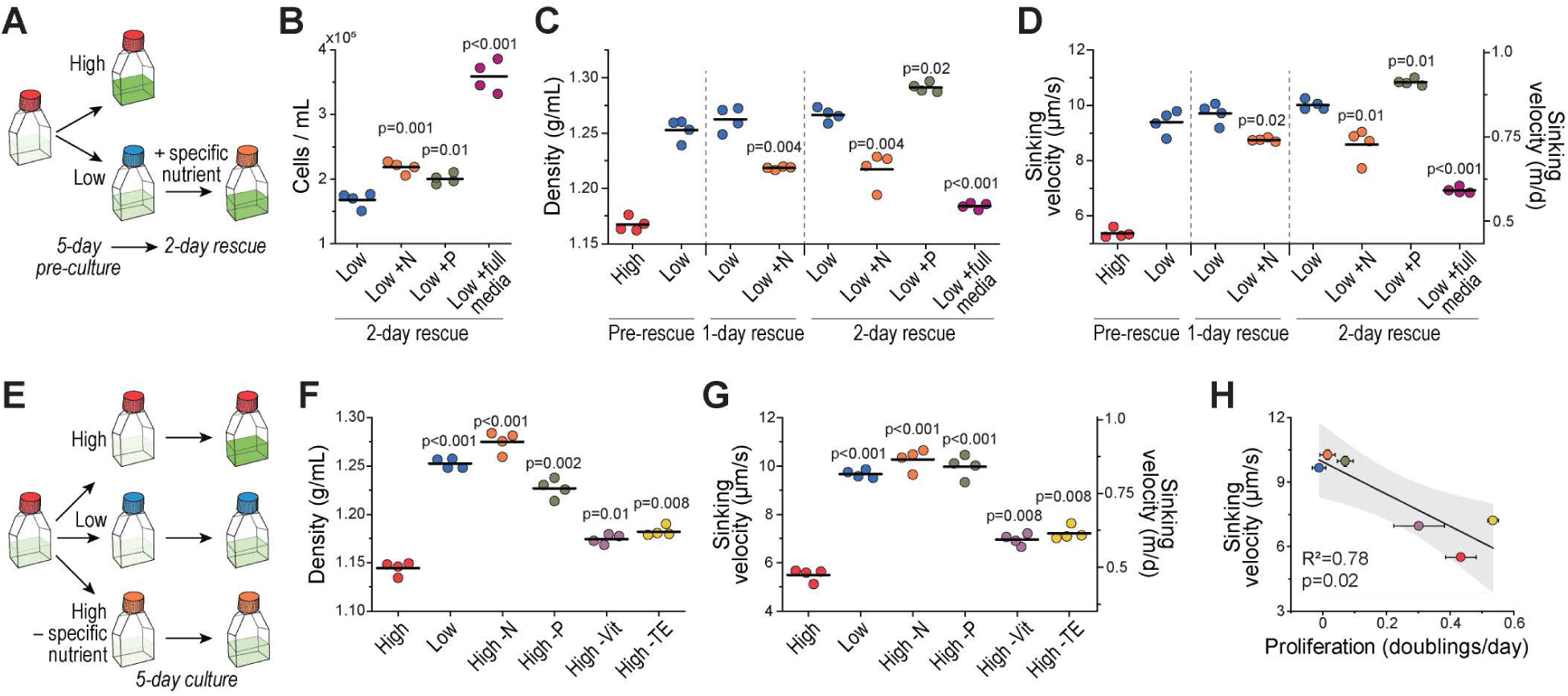
*Tetraselmis sp.* sinking velocity responds to multiple nutrients and correlates with cell proliferation. **(A)** Experimental setup for identifying the limiting nutrient after 5-day culture under low nutrient conditions. **(B)** Cell counts 2 days after addition of indicated nutrient to the low nutrient culture. **(C-D)** *Tetraselmis sp.* cell densities (C) and sinking velocities (D) before and after addition of indicated nutrients to the low nutrient culture. p-values were obtained by paired t-test and depict comparisons to the low nutrient condition from the same day. **(E)** Experimental setup for testing nutrient limitations by specific nutrients. **(F-G)** *Tetraselmis sp.* cell densities (F) and sinking velocities (G) after 5-day culture under indicated nutrient condition. p-values were obtained by paired t-test and depict comparisons to the high nutrient condition. **(H)** Correlation between cell proliferation rate and sinking velocities in the samples used in panel (G). Dots and whiskers depict mean±SEM. Black line and shaded area indicate a linear fit and 95% confidence band. In panels B-G, N indicates nitrogen (orange), P indicates phosphorus (green), Vit indicated vitamins (light purple), and TE indicates trace elements (yellow). Dots depict independent cultures (N=4) and horizontal lines depict mean values.

We then tested if a rescue of the nitrogen limitation can reverse the increased sinking velocities observed under starvation. Supplementation with the complete high nutrient media rescued cell densities and sinking velocities near completely to the pre-starvation levels (Fig. 5C-D), indicating that the sinking velocity changes are reversible. Supplementation of starved cells with additional nitrogen also decreased cell densities and sinking velocities, although this rescue was not complete (Fig. 5C-D). In contrast, supplementation with additional phosphorus increased cell densities and sinking velocities (Fig. 5C-D). This is consistent with previous reports in *Tetraselmis subcordiformis*, where phosphorus increases carbohydrate storages under nitrogen limitation ^34^. Thus, our results indicate that nitrogen limitation is, at least partly, responsible for the increased sinking velocities in starved *Tetraselmis sp*.

Having established that nitrogen limitation can increase sinking velocities, we wanted to test if the regulation of cell sinking velocities is similarly responsive to other nutrient limitations. We cultured *Tetraselmis sp.* under high nutrient conditions where a specific nutrient was depleted (Fig. 5E). After a 5-day culture, limitation of all tested nutrients (nitrogen, phosphorus, vitamins, or trace elements) increased cell densities and sinking velocities in comparison to the complete high nutrient culture conditions (Fig. 5F-G). The sinking velocity increases were driven by cell mass and density increases, rather than cell volume increases (Supplemental Figs. S3A-B), suggesting a similar mechanism of sinking velocity regulation under all conditions. We then correlated the cell proliferation rates across these conditions with the cell sinking velocities. This revealed a strong correlation (R^2^=0.78, Fig. 5H), suggesting that the mechanism regulating cell sinking is coupled to the regulation of cell proliferation, when light is available. Our results indicate that cell starvation, independently of the limiting nutrient, can increase sinking velocities. However, the regulation of size and composition is also impacted by the nutrients that are not growth limiting, as indicated by the observation that phosphorus supplementation can increase sinking velocities further when nitrogen is growth limiting (Fig. 5C-D). We therefore tested additional nutrient conditions by providing *Tetraselmis sp.* with a specific nutrient supplementation at the start of low nutrient culture (Supplemental Fig. S4A). As expected, the early nitrogen and phosphorus supplementation partially rescued cell proliferation (Supplemental Fig. S4B). However, nitrogen supplementation increased cell volumes by ∼45% (Supplemental Fig. S4C), without altering cell densities (Supplemental Fig. S4D). In contrast, phosphorus supplementation had little effect on cell volumes, but increased cell densities (Supplemental Fig. S4C-D). Consequently, both nitrogen and phosphorus supplementation increased cell sinking velocities when supplied pre-starvation (Supplemental Fig. S4E). Altogether, these results indicate that the mechanism regulating cell sinking integrates signals from multiple nutrients.

## DISCUSSION

We have introduced a new approach for measuring gravitational sinking velocities across unicellular algae. Using our new approach, we show that volume is a good predictor of gravitational sinking velocity (R^2^ ∼0.9) when examining different sized species, but not when examining single cells within a species or when comparing the same species under various environmental conditions. This is because cell volume can vary by orders of magnitude between species while cell density cannot (densities outside the range of ∼1.0 – 1.4 g/mL would require extraordinary compositional changes). We also revealed that algae follow the ⅔-power law scaling for cell volume-dependent sinking, both within and between species. This contrasts with a previous meta-analysis, which suggested that larger species sink significantly slower than expected based on the ⅔-power law scaling, possibly due to the extreme cell shapes examined ^31^. Notably, our results are specific to gravitational sinking, whereas previous analyses relied on data from cell settling experiments that are influenced by motility. In motile species, the ⅔-power law scaling of cell sinking may force larger cells to consume more energy to counteract excessive sinking. Because cell size increases in coordination with cell cycle progression ^35–37^, the size-dependent sinking within a species can also result in cell cycle-dependent vertical positioning of the cells in the ocean, as seen in some non-motile species ^15^.

Cell compositional changes are known to impact cells’ vertical movement in non-motile algae ^13–17^. Our work has expanded this concept to motile algae, as we have revealed increased sinking velocities under starvation (stationary phase) due to increased cell densities in motile species. In *Tetraselmis sp.*, we can mechanistically attribute the sinking velocity increases to photosynthesis-driven accumulation of carbohydrates and increased cell density (Fig. 3 and 4). This is consistent with observations in a related species, *Tetraselmis subcordiformis*, where starvation increases starch production ^34^. Overall, our findings support the larger paradigm that uncoupling of photosynthesis and biosynthesis may have important biogeochemical consequences in part by influencing cell sinking ^38,39^.

Based on our observations, we propose a biological mechanism that contributes to the vertical movement of motile single cells in the marine carbon pump and nutrient cycles (Fig. 6). In the photic zone, photosynthesis drives the cellular accumulation of carbohydrates, which increases cell mass and density when nutrients (i.e., nitrogen) are not available to promote cell volume increases and proliferation. This results in faster gravitational sinking, vertically transporting the cells to deeper ocean layers. If the cells encounter nutrients deeper, they can resume proliferation and decrease their gravitational sinking to support vertical upwards movement, as driven by motility and upwelling. *Tetraselmis sp.* can swim faster than the measured sinking velocities ^40^, making upward migration from deeper ocean layers possible, given sufficient energy supply, which the accumulated carbohydrates can supply. Such ability of algae to vertically migrate long (∼100 m) distances has been suggested before ^2,41^, especially for non-motile species ^2,15,16,42^. We therefore propose that the regulation of gravitational sinking in motile algae could impact marine carbon and nutrient fluxes, and this should be accounted for when modeling the marine nutrient and carbon fluxes ^10,11^.

**Fig. 6.**
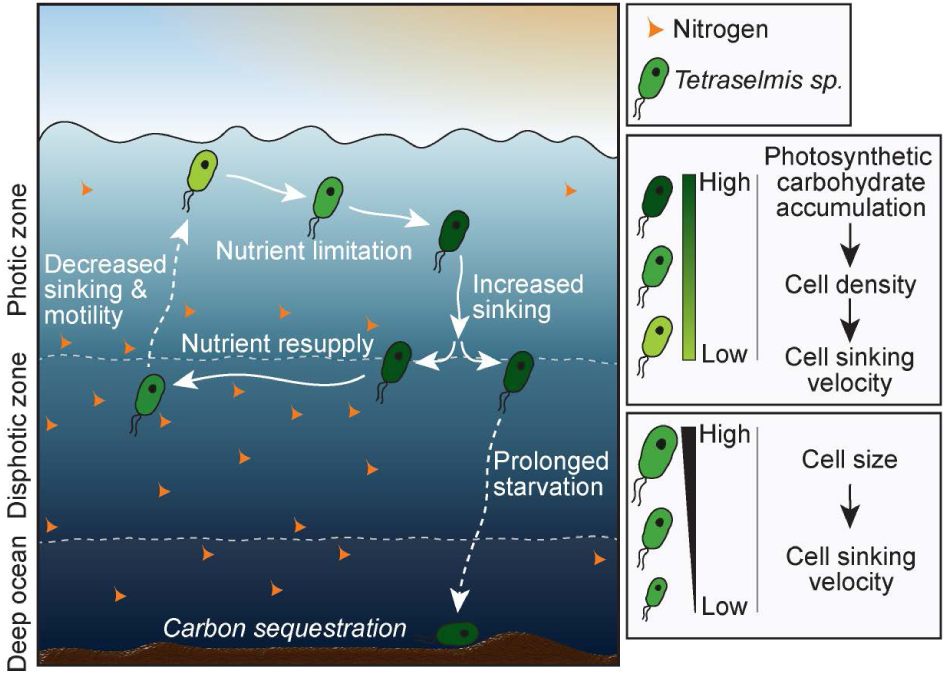
Mechanistic basis for the ecology-dependent vertical movement of unicellular algae. Our results on *Tetraselmis sp.* suggest that the vertical movement of motile algae is partly reliant on the regulation of cell size and composition, which define the cell’s gravitational sinking. In the photic zone, photosynthesis drives the accumulation of carbohydrates, which increases cell mass when other nutrients (i.e., nitrogen) are not available to promote cell volume increase and proliferation. This results in higher cell density and faster gravitational sinking, transporting the cells to deeper ocean layers. If the cells encounter nutrients, they can resume proliferation and decrease their gravitational sinking. The accumulated carbon storages may provide energy for upwards motility. If the cells do not encounter nutrients as they sink, prolonged starvation may result in cell death and eventual sequestration of the cell’s carbon content at the ocean floor. In addition, the vertical cell positioning will also be impacted by cell size, as sinking velocities scales with cell volumes according to the ⅔ power scaling.

Our work also provides further insights into the regulatory mechanism that controls cell sinking. We observed cases where sinking velocity increased due to cell density but not volume increases, and other cases where sinking velocity increased due to cell volume but not density increases. Therefore, the regulation of cell sinking is at least partially independent from the regulation of cell size (volume regulation). This may be because cell volume can impose ecological constraints on the algae by influencing, for example, predation (grazing and viral infection), respiration, and light harvesting ^43–46^, whereas cell density is ecologically less constrained. The regulation of cell sinking can also integrate signals from multiple nutrients, as indicated by different cell sinking responses observed when supplementing the cells with non-growth limiting nutrients. Further understanding of how these signals are integrated, and how they are coupled with cell proliferation, may enable external control of algae sinking that supports carbon capture and biomanufacturing ^47^.

Finally, our work is a proof of methodology that could support research into marine ecology, algae cell biology and the modeling of marine carbon and nutrient fluxes. As exemplified by our findings, our measurement approach is especially valuable in the study of motile unicellular algae whose motility prevents direct live cell investigations of gravitational sinking. While our measurements were carried out in artificial seawater medium and under laboratory conditions, our approach can be utilized to study a variety of algae in aquatic samples. With the development of automated sample preparation methods that separate large algae aggregates from single cells of different sizes, the SMR could be adapted for measurements of algae directly from the ocean. Such method development would enable high throughput sinking velocity measurements *in situ*.

## MATERIALS AND METHODS

### Algal culture conditions

All algae were obtained from Provasoli-Guillard National Center for Marine Algae and Microbiota (NCMA) and belong to the Culture Collection of Marine Phytoplankton (CCMP), except for two coastal seawater isolates (A & B), which were both identified as *Thalassiosira sp.* according to 18S sequencing. Unialgal cultures were grown in filter-sterilized L1 or L1-Si medium, as appropriate for cell type (Table 1). Cultures were maintained at 20°C and they received approximately 50 μm/s/m^2^ of photosynthetically active radiation (PAR) as measured by a LI-COR LI-250 light meter. Cultures were transferred to fresh media weekly by a 1:10 dilution under a laminar flow hood and were regularly visually monitored for health (color, cell shape and size, purity, and motility when relevant) with microscopy.

**Table 1.** List of species studied, their CCMP ID and culture maintenance medium.

| Species | Class | Culture Collection ID | Medium |
| --- | --- | --- | --- |
| *Micromonas pusilla* | Mamelliophyceae | CCMP 1545 | L1-Si |
| *Pyramimonas parkeae* | Prasinophyceae | CCMP 725 | L1-Si |
| *Tetraselmis sp.* | Chlorophyta | CCMP 908 | L1-Si |
| *Chlamydomonas sp.* | Chlorophyta | CCMP 235 | L1-Si |
| *Rhodomonas salina* | Cryptophyta | CCMP 1319 | L1-Si |
| *Bigelowiella natans* | Chloarachniophyta | CCMP 2755 | L1-Si |
| *Nannochloropsis oculata* | Eustigmatophyta | CCMP 525 | L1-Si |
| *Phaeodactylum tricornutum* | Bacillariophyta | CCMP 632 | L1 |
| Isolate A (*Thalassiosira* sp.) | Bacillariophyta | N/A | L1 |
| Isolate B (*Thalassiosira* sp.) | Bacillariophyta | N/A | L1 |
| *Thalassiosira pseudonana* | Bacillariophyta | CCMP 1335 | L1 |
| *Prorocentrum minimum* | Dinophyta | CCMP 1329 | L1-Si |
| *Phaeocystis globosa* | Prymnesiophyceae | CCMP 629 | L1-Si |
| *Emiliana huxleyi* | Prymnesiophyceae | CCMP 2090 | L1-Si |
| *Emiliana huxleyi* | Prymnesiophyceae | RCC 3962 | L1-Si |
| *Isochrysis galbana* | Prymnesiophyceae | CCMP 1323 | L1-Si |
| *Diachronema lutheri* | Pavlovophyceae | CCMP 1325 | L1-Si |

For overall nutrient limitation experiments, two types of media were used: Standard L1 or L1-Si for “high nutrient” conditions, and L1/100 or L1-Si/100 for “low nutrient” conditions, in which the phosphorous, nitrate, trace metals, vitamins, and silica (if applicable) were diluted 1/100, while the salinity and pH of the medium remained unchanged. Algal cultures were pre-grown in maintenance medium for 2 days under the light conditions detailed above. Cells were pelleted by centrifugation at 5000 rpm for 10 min. The supernatant was discarded, and cells were washed once in artificial seawater (pH = 8.2) to prevent nutrient carryover. Cultures were then transferred to the appropriate medium for cell type and desired nutrient availability by a 1:10 dilution under a laminar flow hood and placed in the appropriate lighting condition for 5 days. The three lighting conditions used were “low light”, in which light was available from nearby window at 50 μm/s/m^2^ of PAR and day length varying between 9 h and 15.3 h depending on time of year, “no light” in which cultures were incubated in a drawer without light exposure, and “high light” in which cultures were grown with 100 μm/s/m^2^ of PAR using full spectrum LEDs with a day-night cycle of 11h:13h in a temperature-controlled environmental chamber at 18°C. All data in Figure 1 is from species grown under “low light” and “high nutrient” conditions.

When defining the specific nutrient that limits cell growth and sinking velocity, cells were grown under low nutrient and low light conditions for 5 days, as detailed above, after which the culture medium was supplemented with indicated nutrients to rescue cell growth and/or sinking velocity. The nutrients concentrations used to rescue cell growth and/or sinking velocity were identical to those found in the standard L1-Si media: 8.82 M NaNO_3_, 3.62 M NaH_2_PO_4_H_2_O, L1 vitamin solution, L1 trace element solution, and indicated combinations of these nutrients. The only exception to this was when rescuing cells with “full media” in Figure 3, the low nutrient media was diluted 1:1 with high nutrient media to avoid cell perturbations by centrifugation. The rescue efficiency was determined by cell counting and by OD_600_ measurements 2 days following the nutrient spike-in.

### Phylogeny construction and species verification

Purity and identity of all cultures was verified by Sanger sequencing of the V4 – V9 region of the 18S rRNA gene. To amplify the 18S rRNA gene, cells were lysed by three consecutive freeze-thaw cycles and 1 ul of lysate was added to the New England Biolabs Taq 5X MasterMix. The V4-V9 region was amplified using 570F [5’-CGG TAA YTC CAG CTC YAV] and EukB [5’-TGA TCC TTC TGC AGG TTC ACC TAC] primers added to a final concentration of 0.2μM each. PCR cycling conditions were according to manufacturer specifications, with an annealing temperature of 52°C and extension time of 60 seconds. Purity of the amplification product was verified by gel electrophoresis and PCR products were sequenced by Sanger sequencing at Azenta Life Sciences. The resulting sequences were searched by blastn against the NCBI nonredundant nucleotide collection to verify identy and purity of each strain.

The phylogeny was built by downloading the closest match full-length 18s sequence for each strain from NCBI, along with a representative of land plants, ameobae, and animals (NCBI accession numbers XR_004853735, AF114438, and M10098, respectively). Sequences were aligned using Mafft ^48^ v7.407 default parameters, alignments were trimmed using TrimAL ^49^ v1.2rev59 default parameters, and the tree was inferred using IQ-TREE ^50^ v1.6.8 with the model TIM2e+I+G4 as selected by the IQTree model finder. The resulting phylogeny was visualized and edited with FigTree ^51^ v1.4.4.

### Microscopy-based volume and shape measurements

Cell volume and shape were determined by microscopy of unialgal cultures by EnviroScience, Inc (OH, USA). Live cultures were shipped overnight from Cambridge, MA to Stow, OH at ambient temperature. Upon arrival, 0.1 mL was places into a Palmer-Maloney nano-plankton counting chamber and allowed to settle. In the case that cells are too motile to acquire clear images, the sample was fixed with 1 drop of Lugol’s solution. A microscope-mounted camera was used to acquire images and dimension and volume measurements were acquired by automated image processing software.

### Macromolecular composition analysis

*Tetraselmis sp.* (CCMP 908) was pre-grown in biological triplicates in maintenance medium for 2 days under benchtop light conditions, then transferred by a 1:10 dilution into 100 mL of L1-Si medium for the high nutrient condition and 250 mL of L1-Si/100 medium for the low nutrient condition. After 5 days of incubation under benchtop light conditions, the cultures were pelleted by centrifuging at 10,000 g for 15 minutes at 4°C, resulting in approximately 50 mg of wet biomass per replicate for both conditions. Samples were flash-frozen on a dry ice/ethanol bath and shipped on dry ice to the Bigelow Laboratory for Ocean Sciences (ME, USA), where subsequent analyses were performed by Bigelow Analytical Services. Briefly, dry mass was determined by lyophilizing samples and measuring on an analytical balance. Ash weight was determined as the proportion of initial dry weight remaining after the sample was placed in a muffle oven at 600°C for 16 hours. The proportion of protein in the dry mass was determined via the Dumas Method using a Costech ECS 4010 Elemental Analyzer utilizing a Nitrogen-to-Protein conversion factor of 4.76 ^52^. The proportion of lipid in the dry mass was determined by an amended GC/MS method ^53^. Individual fatty acids were measured, then all fatty acids were summed. The proportion of carbohydrates was determined indirectly as the remaining mass after accounting for water, ash, protein, and lipids.

### Cell count, buoyant mass, and volume measurements

Buoyant mass measurements were carried out using a suspended microchannel resonator (SMR). The SMR is a vibrating cantilever with an internal microfluidic channel that allows single cells to travel through the interior of the cantilever. The resonant frequency of the cantilever changes due to the presence of a cell, and this change is proportional to buoyant mass of the cell. The fabrication and details of the operation of SMR are detailed in ^19,22,54,55^. This study used two different sizes of SMRs, one where the cantilever inner cross-section is 8 µm x 8 µm, and one where the cross-section is 15 µm x 20 µm. Algae species with estimated diameter below 6 µm were measured using the smaller SMR, and larger algae species were measured using the larger SMR. For each measurement, an aliquot of cell culture was transferred to a vial fluidically connected to the SMR, and single cells were flushed through the SMR in culture media. Each cell measurement lasted <100 ms (travel time through the cantilever). All measurements were carried out between 10 am and 6 pm.

Cell volumes were measured using a Coulter counter (Multisizer 4, Beckman Coulter), which is based on the principle that a cell traversing an orifice filled with electrolyte will produce an impedance change proportional to the volume of the cell due to displacement of electrolyte. Algae cells were measured with one of two Coulter counter apertures, one 20 µm and one 100 µm in diameter, with the selection of aperture depending on cell size. Cell counting was also based on coulter counter measurements. Each cell is measured in ∼1 ms (travel time through the coulter counter aperture). All measurements were carried out in culture media. The background noise of the SMR and coulter counter measurements was determined using a blank sample (no cells), and particles smaller than the reliable detection limit were not analyzed. When analyzing cell proliferation rates, cell counts were collected on days 4, 5, and 6 to calculate proliferation rate assuming exponential growth (sinking velocity measurements were carried out on day 5).

For SMR data analysis, the real-time resonant frequency data were analyzed by custom MATLAB code which detects the resonant frequency change caused by passage of a cell ^55^. The frequency change for each cell was converted to buoyant mass of the cell based on calibration using NIST-certified polystyrene beads (3 µm and 10 µm in diameter, Duke Standards, Thermo Scientific). For Coulter counter data analysis, cell volumes are automatically available from the instrument, and these values were re-calibrated based on measurements of NIST-certified polystyrene beads (2 µm and 6 µm in diameter, Duke Standards, Thermo Scientific).

### Sinking velocity calculations

Algae sinking velocities were calculated according to the Stokes’ law (*equation 1*), using the following values for environmental constants in surface ocean: the dynamic viscosity of seawater of 1.07 × 10^-3^ Pa·s at 20°C, and density of seawater of 1.02 g/ml. Both high- and low-nutrient culture medias displayed similar densities, as quantified by SMR following a previously reported protocol ^20,56^. Cell radius was calculated from cell volume and cell density was calculated using the volume and buoyant mass measurements (*equation 2*). In all agal species tested in this study, the Reynolds numbers did not exceed the scale of 10^-7^.

The Stokes’ law drag force was corrected to account for non-spherical cell shapes. The correction was carried out according to previous empirical results for a variety of shapes ^18^. The shape and aspect ratio for each species were first determined based on microscopic images obtained in this study and/or provided by algae supplier (NCMA). A multiplicative shape factor was calculated for each species based on its shape and aspect ratio. All results shown in figures are without shape correction, as the calculated corrections were too minor to be clearly visible on the graphs (except for a single species, *Phaeodactylum tricornutum*).

### Single-cell density and sinking velocity measurements

Single-cell densities and sinking velocities were obtained by coupling buoyant mass measurements with fluorescence exclusion-based volume measurements ^24^ within the SMR device by building a fluorescence microscope on top of the SMR ^23^. In fluorescence exclusion measurements, the media contains a fluorophore, which is cell impermeable. When the fluorescence signal from a fixed volume is measured with and without a cell, the signal intensity decreases according to the volume of the cell as fluorescent media is excluded from the measurement volume. In practice, cells were immersed in culture media containing 5 mg/mL Fluorescein isothiocyanate–dextran (Sigma-Aldrich, Cat#FD2000S, molecular weight 2,000,000 g/mol) and flown through the SMR. Immediately after the SMR cantilever, the cells flowed through a fluorescence measurement area where the sample was exited with 482 nm light. Emission from the fluorescent dextran was collected with a photomultiplier tube using 515/30 nm emission filter. A typical measurement duration is 2 ms (time that the cell travels through the fluorescence measurement area).

SMR and fluorescence exclusion measurement data were paired using time information for each cell measurement, with a typical time lag between the two measurements being 10 ms. Cell volume was obtained by calculating the ratio between fluorescence signal decrease when a cell travels through the measurement area and the fluorescence baseline when a cell is not present in the measurement area. These cell volumes were then calibrated to absolute volume units by comparing the population size distributions to those measured with Coulter Counter. Single-cell densities and sinking velocities were calculated from the buoyant mass and volume data as detailed above for population-level measurements.

### Data analysis and statistics

In all figures, n refers to the number of single cells measured, while N refers to the number of independent experiments. Particles too small to be viable cells were removed from all final analyses. For single-cell density measurements, particles with density above 1.7 g/mL were excluded, as these were likely due to mispairing of mass and volume measurements. For analyses of volume-dependency, data was log-transformed and then fitted using a linear regression. When comparing volume-dependency across species, reported errors for scaling exponent reflect the error of the fitted slope. When comparing volume-dependency among single cells within a species, reported errors for scaling exponents reflect the error between independent experiments. Phylogenetically informed regression was performed by fitting a linear model to density, log-transformed volume, or log-transformed sinking velocity and including the phylogeny as a covariate using the phylolm package ^57^ in R (2021). Pagel’s lambda model was used for testing phylogeny as a covariate in our data. All other statistical details indicated in the figures and figure legends and details were calculated using OriginPro 2023 software.

### Data availability

All buoyant mass and volume data, which were used to calculate cell densities and sinking velocities, are included in the Supplemental datasets S1-S10. For details about the Supplemental datasets, please see the first datasheet (readme) in the supplemental datafile.

## Author contributions

T.P.M., A.L.G., Y.W., M.F.P. and S.R.M. conceived and designed the research; T.P.M., A.L.G., Y.W., W.W., T.R.U. and Y.H. carried out the experiments; T.P.M., A.L.G., Y.W. and B.R.K.R. analyzed the data; B.R.K.R., M.F.P. and S.R.M. provided resources and conceptual feedback; and T.P.M., A.L.G. and Y.W. wrote the paper with input from all authors.

