## Supplemental Figures for "Direct quantification of unicellular algae sinking velocities reveals cell size, light, and nutrient-dependence"

### SUPPLEMENTAL FIGURES & FIGURE LEGENDS

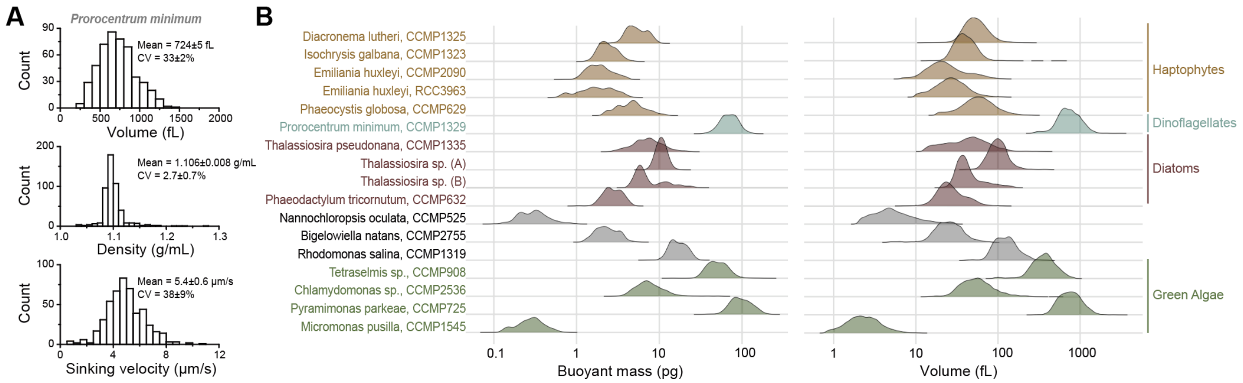


**Supplemental Fig. 1. Cell masses and volumes across phylogenetically diverse algae species.** **(A)** Representative single-cell volume, density, and sinking velocity histograms for *Prorocentrum minimum* (n=484 single cells). The listed statistics depict mean±SD and were calculated by comparing 2 independent experiments. **(B)** Probability density functions of single-cell buoyant masses (left) and volumes (right) of the species indicated on the left. For each species, data is shown for a representative example from multiple independent cultures. Data is the same as used for Fig. 1D.

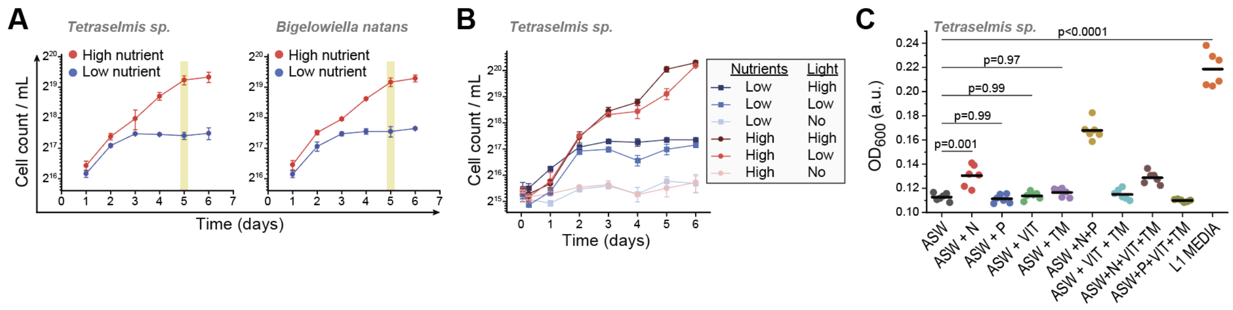


**Supplemental Fig. 2. Nutrient and light-dependency of *Tetraselmis sp.* and *Bigelowiella natans* proliferation.**

**(A)** Cell counts over 6 days for *Tetraselmis sp.* and *Bigelowiella natans* grown under high and low nutrient conditions. Data depicts mean±SD of 4 independent cultures. **(B)** Cell counts over 6 days for *Tetraselmis sp.* grown under indicated nutrient and light conditions. Data depicts mean±SD of 3 independent cultures. **(C)** $OD_{600}$ measurements of *Tetraselmis sp.* culture that were starved under low nutrient conditions for 5 days and then supplemented with indicated nutrients for 2 days. ASW = artificial seawater (no nutrients), N = nitrogen, P = phosphorus, VIT = vitamins, TM = trace metals, L1 media = high nutrient condition growth media (L1-Si). Dots depict independent cultures (N=6 for each condition) and horizontal lines depict mean values. p-values were obtained by ANOVA followed by Tukey's posthoc test.

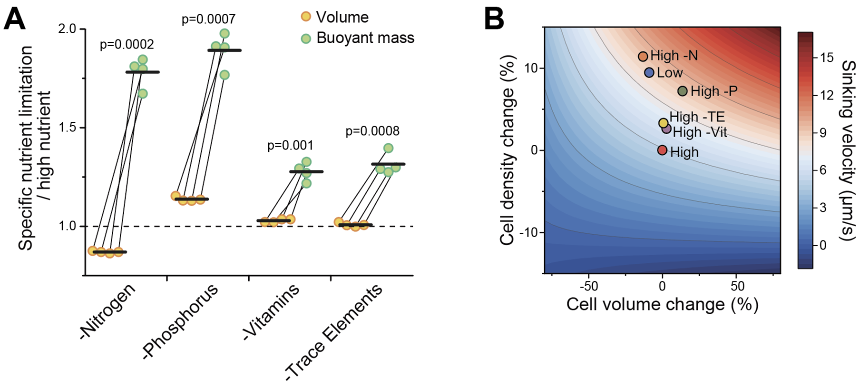


**Supplemental Fig. 3. Various nutrient limitations cause *Tetraselmis sp.* to increase sinking velocity by increasing cell density.**

**(A)** Change in cell volume (orange) and buoyant mass (green) between nutrient conditions that are limited by a specific nutrient (e.g. high nutrient minus nitrogen) and high nutrient conditions. Data is the same as in Fig. 5F-G. Each dot depicts an independent culture (N=4), paired samples are connected by a line, and horizontal lines depict mean values. p-values were obtained by pair-sample t-test and reflect the difference between buoyant mass and volume changes. **(B)** Relative change in cell volume vs relative change in cell density upon indicated nutrient limitations (colored dots). All values are in comparison to the high nutrient condition (red dot). Data and dot color coding is the same as in Figs. 5F-H, and each dot represents the average of 4 independent cultures. The color gradient in the background indicates sinking velocities. The observed cell volume changes have little impact on sinking velocities in comparison to the observed cell density changes.

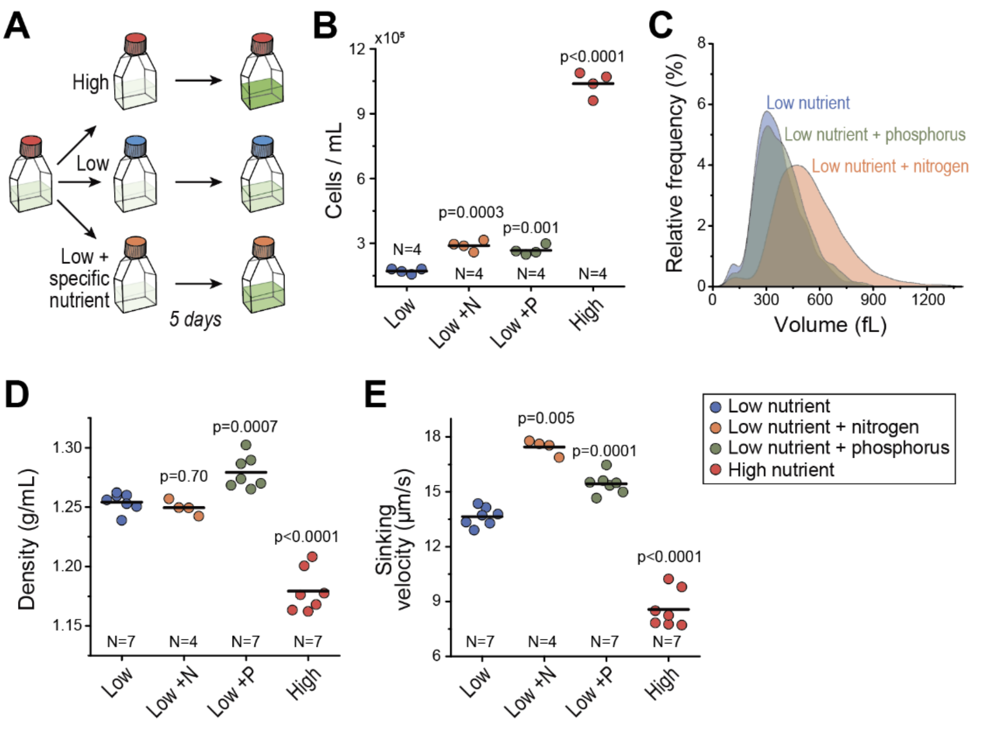


**Supplemental Fig. 4. Nitrogen supplementation to proliferating *Tetraselmis sp.* increases cell size, causing increased cell sinking velocity.**

**(A)** Experimental setup for testing how specific nutrient supplementation of low nutrient media impacts cell size, density and sinking velocities. **(B)** *Tetraselmis sp.* cell counts after 5-day culture under indicated nutrient conditions. N indicates nitrogen (orange), P indicates phosphorus (green). **(C)** Probability density functions of *Tetraselmis sp.* single-cell volumes after 5-day culture under indicated nutrient conditions. Nitrogen supplementation increased cell volume significantly ($p<0.0001$, paired t-test, N=4 independent experiments). **(D-E)** *Tetraselmis sp.* average cell densities (D) and sinking velocities (E) after 5-day culture in indicated nutrient conditions.

In panels (B), (D) and (E), dots depict independent cultures, N depicts the number of independent cultures and horizontal lines depict mean values. p-values were obtained by paired t-test and depict comparisons to the low nutrient condition.
